# Estimation of effective concentrations enforced by complex linker architectures from conformational ensembles

**DOI:** 10.1101/2021.08.13.456233

**Authors:** Magnus Kjaergaard

**Affiliations:** Department of Molecular Biology and Genetics, Aarhus University; The Danish Research Institute for Translational Neuroscience (DANDRITE), Nordic EMBL Partnership for Molecular Medicine; Center for Proteins in Memory - PROMEMO, Danish National Research Foundation

## Abstract

Proteins and protein assemblies often tether interaction partners to strengthen interactions, to regulate activity through auto-inhibition or -activation, or to boost enzyme catalysis. Tethered reactions are regulated by the architecture of the tether, which defines an effective concentration of the interactor. Effective concentrations can be estimated theoretically for simple linkers via polymer models, but there is currently no general method for estimating effective concentrations for complex linker architectures consisting of both flexible and folded domains. We describe how effective concentrations can be estimated computationally for any protein linker architecture by defining a realistic conformational ensemble. We benchmark against prediction from a worm-like chain and values measured by competition experiments and find minor differences likely due to excluded volume effects. Systematic variation of the properties of flexible and folded segments show that the effective concentration is mainly determined by the combination of the total length of flexible segments and the distance between the termini of the folded domains. We show that a folded domain in a disordered linker can increase the effective concentration beyond what can be achieved by a fully disordered linker by focusing the end-to-end distance at the appropriate spacing. This suggests that complex linker architecture may have advantages over simple flexible linkers and emphasizes that annotation as a linker should depend on the molecular context.

## Introduction

Many biochemical reactions occur internally in molecules or protein complexes including auto-inhibition, multivalent binding, or tethered enzyme reactions. When the reactants are physically connected, reactions are first-order and reaction half-life is thus independent of concentration. Instead, reaction rates depend on how the interacting entities are connected, which enables allosteric regulation of tethered reactions via the linkers.^1^ The prototypical tether is a flexible linker in the form of an intrinsically disordered region (IDR), a protein region that does not have a fixed tertiary structure but is active as a disordered chain.^2^ Disordered linkers are ubiquitous in multidomain proteins, and functional annotations suggest that “linker” is the most common function for IDRs^3^. Furthermore, linkers play an important role in biotechnology, where they tether domains to form multivalent binders,^4^ multi-enzyme cascades^5^ or genetically encoded sensors and switches.^6^ In such proteins, optimizing the linker is the crucial step for tuning activity. To understand structure-function relationships for linkers, it is necessary to understand how the properties of the linker determine the contact probability of the entities they link.

Molecular tethers are often more complex than a continuous flexible region. In the auto-phosphorylation of calmodulin-dependent kinase II, the catalytic domain phosphorylates a regulatory region in a neighboring subunit. The tether controlling this reaction consists of an IDR, the hub domain of the kinase, and another IDR.^7^ In the bivalent interaction between the PDZ domains of PSD-95 and ionotropic glutamate receptors, the tether consists of the disordered C-terminal domain of one receptor subunit, the ion channel domain, and the disordered C-terminal tail of another receptor subunit.^8^ The connections that tether biochemical reactions thus vary beyond what is typically considered linkers and may include domains that have other functions. In the following, the term “linker architecture” will be used to encompass complex connections that may consist of several rigid and flexible segments, and the term “linker” will be reserved for a single continuous IDR.

Linker architectures enforce an effective concentration (*C_eff_*) of the tethered ligand, which corresponds to the concentration of free ligand that has the same encounter rate as the tethered ligand. When the effective concentration is known, it is possible to describe a tethered system via the equilibrium and rate constants of the corresponding untethered system. Quantitative models of *C_eff_* has been used to describe processes such as auto-inhibition,^9,10^ avidity of multivalent interactions,^11–17^ phase separation^18^ and intra-complex enzyme catalysis.^19^ In principle, *C_eff_* can be measured directly in competition experiments, where a free ligand displaces the tethered ligand.^20,21^ Such competition experiments require an experimental read-out of intramolecular binding for example via a fluorescent competitor^20^ or a designed FRET biosensor.^21,22^ Direct measurement through competition experiments requires reconstitution of the system *in vitro* and displacement of the intra-molecular ligand, which typically requires millimolar concentrations of competitor. Therefore, experimental determination of *C_eff_* is often impractical, and it is attractive to model *C_eff_* theoretically instead.^11,12,17,23–25^

*C_eff_* is determined by the end-to-end distance distribution of the linker, which defines the probability of a tethered ligand contacting a binding site at a certain spacing from the origin of the linker. Fully disordered linkers can often be modelled well by homopolymer models.^26–28^ Though homopolymer models ignore many aspects of IDRs, they provide a good first approximation, and models such as the worm-like chain (WLC) are currently widely used to predict *C_eff_*. *C_eff_* defined by the continuous end-to-end distance probability as follows:^11,29^

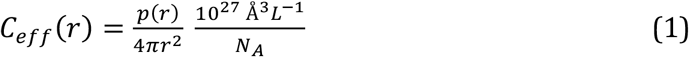

The first term divides the probability distribution by the surface area of a sphere of radius *r*, and the second term converts to molar concentration. As the WLC can be expressed purely mathematically, *C_eff_* enforced by flexible linkers can be calculated rapidly for example by the interactive web server “C_eff_ calculator”.^30^ Purely mathematical descriptions, however, struggle with complex linker architectures composed of e.g. both folded and disordered regions. Simplifying assumptions are thus necessary to handle complex linker architectures. For example, it has been suggested that a folded domain in a WLC can be approximated by rigid spacer with a length corresponding to the distance between the attachment sites irrespective of its position in the linker.^25^ Such descriptions do not account for e.g. excluded volume, and are difficult to validate without direct experimental comparison. This suggests a potential for improvement of the theoretical prediction of *C_eff_* enforced by complex linker architectures.

This work seeks to develop a simple approach for estimating *C_eff_* for complex linker architectures composed of disordered and folded segments. Like the WLC, we aim for method that describes the general geometric principles of a complex linker architectures, but do not account for sequence specific interactions. *C_eff_* can be calculated at different spacings for any linker architecture where we can define a valid end-to-end distribution, although this has not been used to study protein linkers to the best of our knowledge. We thus aim to simulate a realistic structural ensemble for the linker architecture, which allows determination of the distribution of end-to-end distances and *C_eff_* as a function of spacing. Results are compared to results from competition experiments and predictions from the WLC, and subsequently we explore how *C_eff_* depends on the properties of the folded and flexible segments in complex linker architectures.

## Materials and methods

### Ensemble generation

Coarse-grained ensembles that represent each residue as a bead were generated using the RANCH routine of EOM2.0.^31,32^ Unless indicated otherwise, the “native chain” setting was used for flexible segments, and the ensemble size was set at 10.000. Input sequences were generated by adding flanking GS-repeats to the sequences of the folded domains. The second disordered segment either started with G or S as needed to disambiguate pseudo-atoms. For the linkers used in the biosensors,^21^ the residues corresponding to the flanking restriction sites in the corresponding DNA construct were also included. Poly-alanine α-helices were built using PyMOL (v. 1.8.0.5, Schrödinger Inc.) with lengths ranging from 7 to 77 residues. Folded domains were all crystal structures with a single chain without missing residues. For models with several copies defined in the asymmetric unit, the first model was selected. The sequences used by EOM were truncated to the region defined in the PDB model, and co-factors were removed. The structured domains were: 1IIU,^33^ 1MPD,^34^ 1N2A,^35^ 1TM1^36^ (chains E and I separately), 1W6X,^37^ 2GI9,^38^ 2IC2, 2R4U,^39^ 3D7S,^40^ 3GV9,^41^ 3H1C,^42^ 3UC7,^43^ 3MYC,^44^ 4F5S,^45^ 4KDS,^46^ 4O75,^47^ 5D13, 5DFT, 5EJ6.^48^ The r_NC_ distance was measured between the first and last C«-atoms using PyMOL. For the dimeric a-actinin (PDB:1HCI),^49^ the disordered segments were attached to the C-terminus of the folded domain, and distances were measured between the two C-termini.

### Determination of C_eff_

The distance distribution was measured between the first and last bead in each model using a PyMOL script. The resulting list of distances was imported into Rstudio, and a histogram of end-to-end distances was generated in 1 Å bins. The radial probability distribution p(r) is defined as the fraction of conformers with end-to-end distances within 0.5 Å of the bin center. For each bin, p(r) and r were inserted into (Eq. 3) to calculate *C_eff_* For the WLC, the p(r) function was predicted using the “*C_eff_* calculator” web applet (http://Ceffapp.chemeslab.org)^30^ and *C_eff_* was calculated for each point using (Eq. 1).

## Results

We aimed to develop a simple strategy for estimating *C_eff_* enforced by complex linker architectures such as mixtures of folded and disordered segments. We considered the case where a tethered ligand binds a site located at a fixed spacing from the origin of the linker (Fig. 1A). An analogous situation occurs for bivalent interactions after binding of the first ligand, which has been the focus of previous studies of linker architecture due to its importance in drug development. We made the simplifying assumption that the linker architecture can be divided into flexible linkers and rigid domains (Fig 1B). Ensembles were generated by the random chain algorithm of the Ensemble Optimization Method (EOM)^31,32^, which models flexible linkers as self-avoiding chains sampling backbone dihedral angles from one of three different coil libraries (Fig. 1C). EOM was originally developed to reproduce small angle x-ray scattering data of flexible proteins and thus allows inclusion of folded domains as rigid bodies, which is the function needed here. EOM also has a built-in genetic algorithm that allows selection of an ensemble that agrees with experimental data. This function will not be used here, where we just use the ensemble based on the coil-library. End-to-end distances were extracted from the ensemble and counted in 1 Å bins, which can be visualized as a probability density distribution histogram (Fig. 1D). For each combination of r and *p(r), C_eff_* can be calculated as described below, and visualized as *C_eff_* at different spacings from the attachment site (Fig. 1E), which will hereafter be referred to as the *C_eff_* profile.

**Figure 1:**
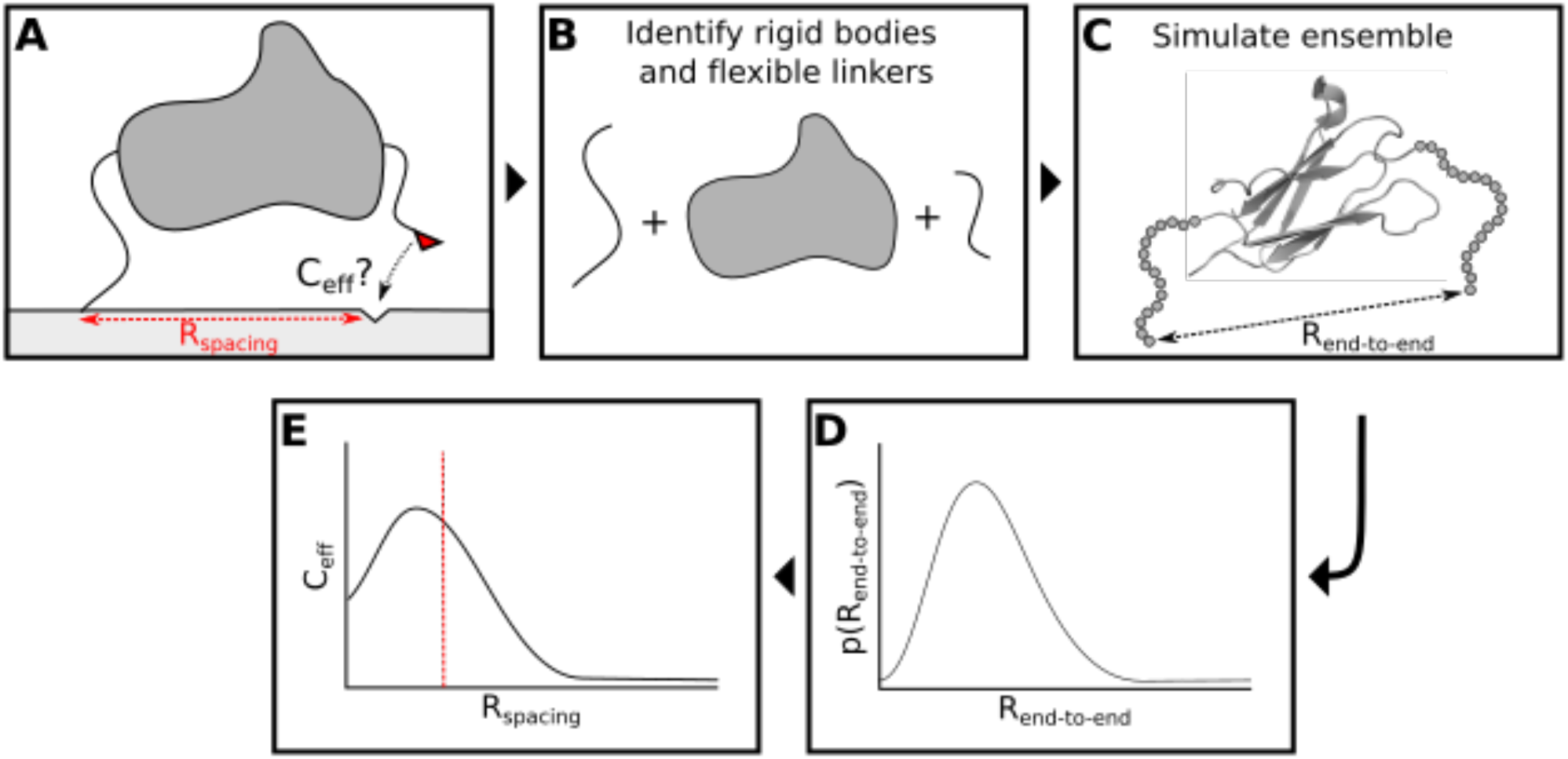
Flowchart for the estimation of C_eff_ for linker architectures composed of folded and disordered segments. First, the linker architecture was divided into parts that will be treated as rigid bodies and parts that will be treated as disordered chains (B). An ensemble of typically 10.000 conformations was generated for each linker architecture (C). A histogram of end-to-end distances was constructed (D) and converted into C_eff_ of tethered ligand at different spacings from the initial attachment site (E).

### Calculation of C_eff_ from a conformational ensemble

We would like to be able to estimate C_eff_ from a conformational ensemble. Eq. 1 was developed for mathematical polymer models, where the probability distribution is precise and continuous. In a discrete conformational ensemble, no conformers will have an end-to-end distance of precisely 42 Å. Instead, it is practical to consider the probability distribution of end-to-end distances in bins of a certain size such that all conformers with an end-to-end distance between 41.5 and 42.5 A are counted in the 42 A bin. Analogous to Eq. 1, we define *C_eff_* as the ratio of the molar concentration of tethered ligand in each bin, and the volume of the bin:

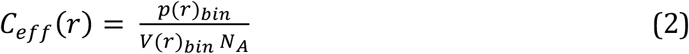

For a 1 A bin centered at 42 Å, *p(r)bin* is simply the fraction of conformers where the end-to-end distance falls between 41.5 and 42.5 Å. *V(r)bin* is the difference in volume between sphere of radii 42.5 A and 41.5 Å. Eq. 3 thus estimates C_eff_ from a discrete conformational ensemble for a generalized bin width of *bw* including a term to convert to units of from Å^3^ to L:

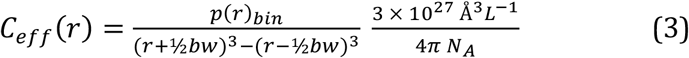

#### Comparison to WLC

We compared the predictions from the EOM ensemble for a fully disordered chain to the worm-like chain. We created ensembles of 10.000 models for (GS)_50_ using the three different coil libraries built into EOM: “Compact”, “Native” and “Random”. Each ensemble can be generated in minutes on a personal computer. The probability distributions were fitted to the WLC model resulting in persistence lengths of 5.2, 6.4 and 7.6 Å, respectively (Fig. 2A-C). The ensemble-derived *C_eff_* profiles were noisy at low spacings at an ensemble size of 10.000, which is likely to be inherent to *C_eff_* determination from ensembles: The close spacings are only represented by few conformers in the ensemble, and values vary stochastically. The noise can be reduced by increasing the ensemble size although at the expense of computational cost, or by increasing the bin size at the expense of spatial resolution.

**Figure 2:**
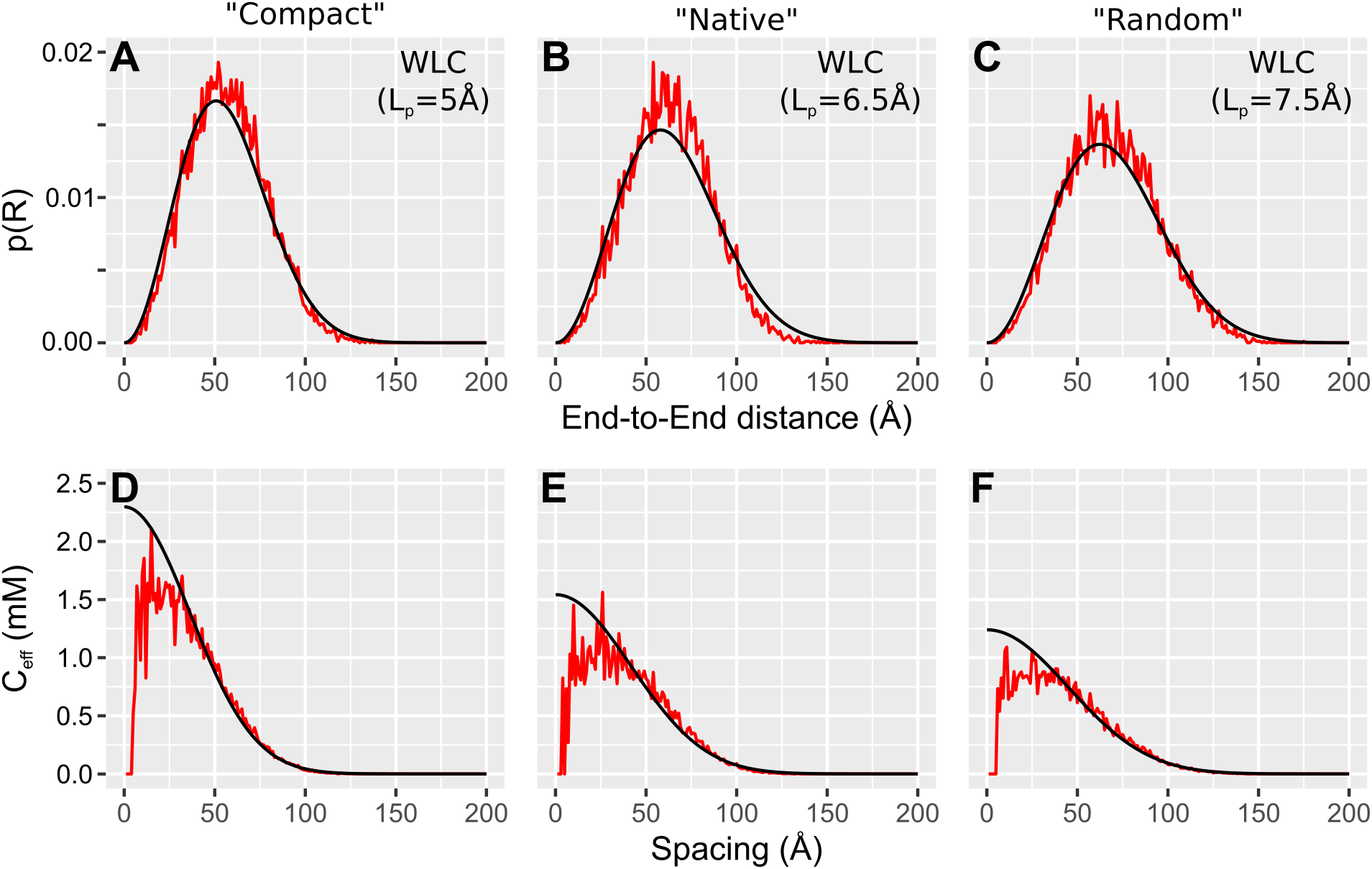
Comparison of ensemble reconstruction and worm-like chain modelling for a fully disordered linker. Ensembles of 10.000 conformations of a (GS)_50_-linker were constructed using the (A, D) “Compact”, (B, E) “Native” and (C,F) “Random” coil libraries of EOM.^12^ WLC simulations were done using “C_eff_ calculator”, using persistence lengths from a fit of the WLC to the p(R), which gave the following values: “Compact”: 5.2 Å (5.09 - 5.30 Å). “Native”: 6.4 Å (6.27 - 6.63 Å). “Random”: 7.6 Å (7.38-7.73 Å). The black lines correspond to from the WLC, and the red line the ensemble. Values in parenthesis represent 95% C.I. of the fitted parameters.

For the remainder of the manuscript, we use ensembles of 10.000 as a practical compromise between speed and precision. *C_eff_* profiles calculated from ensembles and the WLC agreed at spacings above ~20 A (Fig. 2D-F). For shorter spacings, however, the ensemble derived *C_eff_* peaked at what appeared as a broad plateau before decreasing when it approached zero. The *C_eff_* calculated from the WLC increased monotonously to a maximum at zero as shown previously.^29^ Due to the division by r^2^ in Eq. (1), the *C_eff_* profile magnifies differences at small spacings that are easily overlooked in the distance distribution. The different *C_eff_* profile is likely due to differences in how the models encodes self-avoidance. The close spacings report on the subset of the ensemble where the ends of the linkers are close to each other, which increase the importance of steric occlusion. In total, the ensemble derived *C_eff_* values quantitative agree with the WLC, which accurately model many disordered regions.

#### Comparison of length scaling and experimental values

Next, we wanted to compare the ensemble *C_eff_* to values determined from competition experiments using a FRET biosensor.^21^ Experimental *C_eff_* values from the biosensor scaled with linker length according to a polymer scaling law of the following form:^21^

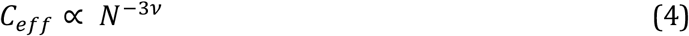

Where *N* is the number of disordered residues and *v* is the exponent of an analogous scaling law for the end-to-end distance. The physical properties of the linkers affect the scaling via the linker compaction, where compact and extended linkers have low and higher values of *v*, respectively. EOM does not account for electrostatics which is the dominant factor in IDP compaction, so we only compared the ensemble values to biosensor containing (GS)_x_-linkers. For each (GS)_x_-linker, we generated 5 ensembles of 10.000 structures and calculated the resulting *C_eff_* at a spacing of 10 Å (Fig. 3), which is the distance between linker attachment sites in the complex used in the biosensor.^50^ The spacing of 10 A is near the noisy maxima in the *C_eff_* profile. While the ensemble size defines the shape of the *C_eff_* profile, individual bins had highly variable *C_eff_* values, and we thus averaged over 3 Å. The competition experiment produced values that are consistently 2-3-fold lower than predicted by the ensemble and the WLC. It is unclear whether this difference represents a systematic error in the prediction or experiment (see discussion).

**Figure 3:**
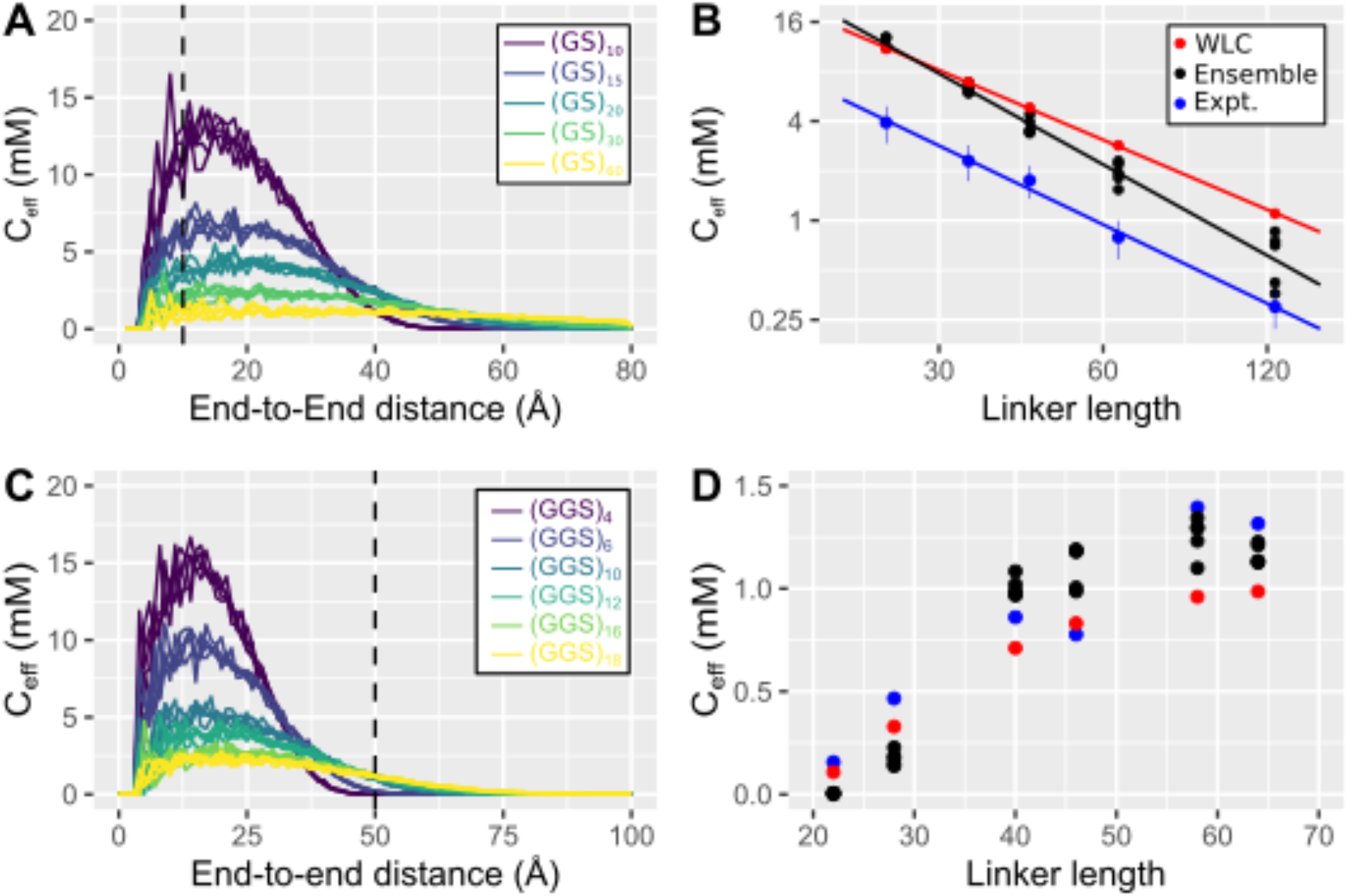
Comparison of predicted C_eff_ for disordered linkers to values measured using FRET biosensors. (A) Effective concentration landscapes predicted for a series of disordered (GS)_x_-linkers previously been measured by competition experiments in a FRET biosensor.^25^ In addition to the (GS)_x_-repeat, the linker contains two residues from restriction sites in either end. For each linker, five ensembles of 10.000 conformations were simulated. The dashed line indicates the spacing (10 Å) between the attachment sites in the complex used in the biosensor.^50^ (B) Linker length scaling of C_eff_ at 10 Å from prediction and experiments. C_eff_ values from ensemble calculation is averages of three 1 Å bins from 8.5-11.5 Å. Scaling exponents from the fits are WLC (L_p_ = 6.4Å): −1.41, ensemble: −1.83 and experiment: −1,59). (C) Effective concentration landscapes predicted for a series of disordered (GGS)_x_-linker used in a Zn^2+^-biosensor, where the attachment sites are spaced by 50 Å (dashed line) in the complex. The linkers contain 10 flexible residues in addition to the GGS-repeat. (D) Comparison of C_eff_ derived from experiment, EOM ensembles and a worm-like chain with an L_p_ of 4.5 Å as determined by fitting to the experimental data

The predicted *C_eff_* depended on linker length via a power law as indicated by the linear relation in the double logarithmic plot (Fig. 3B) in agreement with both experiments and predictions from WLC. Ensemble *C_eff_* values decreased more rapidly with linker length than both the experiments and the WLC (Fig. 3B). The experimental *C_eff_* values for GS-linkers have a scaling exponent of −1.53, which correspond to a scaling exponent for chain length of ν = 0.51. The slight difference from the value reported previously^21^ depends on whether you include the 4 residues from the restriction sites of the corresponding DNA construct. The corresponding values are - 1.40 (v = 0.47) for the WLC and −1.83 (ν = 0.61) for the ensemble. The spacing from the biosensor (10 Å) corresponds to a region, where a systematic deviation was seen between the EOM ensemble and the WLC (Fig. 2), and the difference in scaling exponent was thus expected. Consistent with excluded volume causing this deviation, the scaling exponents from the ensemble depended on the spacing, and a scaling exponent of −1.63 (ν = 0.54) was observed at a spacing of 20 Å. Other biophysical experiments show that IDRs have average v of ~0.50-0.53^28,51,52^, but with a lot of individual variation as highly charged IDRs can have v up to 0.7.^27^ The scaling of *C_eff_* values from the EOM ensemble with linker length corresponded approximately to that expected for an IDR of average compaction. When restricted to an end-to-end distance of 10 Å, excluded volume effects will likely increase for longer chains, which both explains why the scaling exponent depends on the spacing and why predictions differ between the WLC and the ensemble.

We thus tested how the ensemble method performed for longer spacings between the attachment sites where steric effects are not as dominant. Therefore we simulated the linkers previously used to tune a Zn^2+^-biosensor,^53^ where C_eff_ was determined by comparison of titrations of an untethered and a tethered Zn^2+^-dependent interaction for six of GGS-repeat linkers. The spacing between linker attachment sites was 50 Å,^54^ which is longer than the typical end-to-end distance of the shortest linkers and therefore C_eff_ increased with linker length (Fig 3C). We simulated this linker series using EOM and compared it to experimental values and predictions from a worm-like chain with a persistence length determined by fitting the experimental data (4.5 Å). For this linker set, EOM reproduced the shape and magnitude of the experimental C_eff_ values approximately equally well as the WLC (Fig. 3D), but without the need for fitting the persistence length.

#### Folded domains in disordered linkers

Having validated the ensemble method, we wanted to investigate linker architectures that are difficult to describe mathematically. The simplest and most general type of complex linker architecture is a folded domain flanked by two flexible linkers. Mathematical treatment suggests the length of the two disordered segments can simply be added together and considered as a single long linker.^25^ To test this assertion, we generated ensembles of disordered linkers with three different folded domains embedded: A SH3 domain, a PDZ domain and a FN3 domain, which are typical small globular domains found in multidomain proteins. We tested the distance distribution and *C_eff_* profile with a total of 100 disordered residues (Fig. 4A-F), where the flexible segments sample a much larger volume than the folded domains, and 30 residues where they sample a volume similar to the dimensions of the folded domain (Fig. 4G-L).

**Figure 4:**
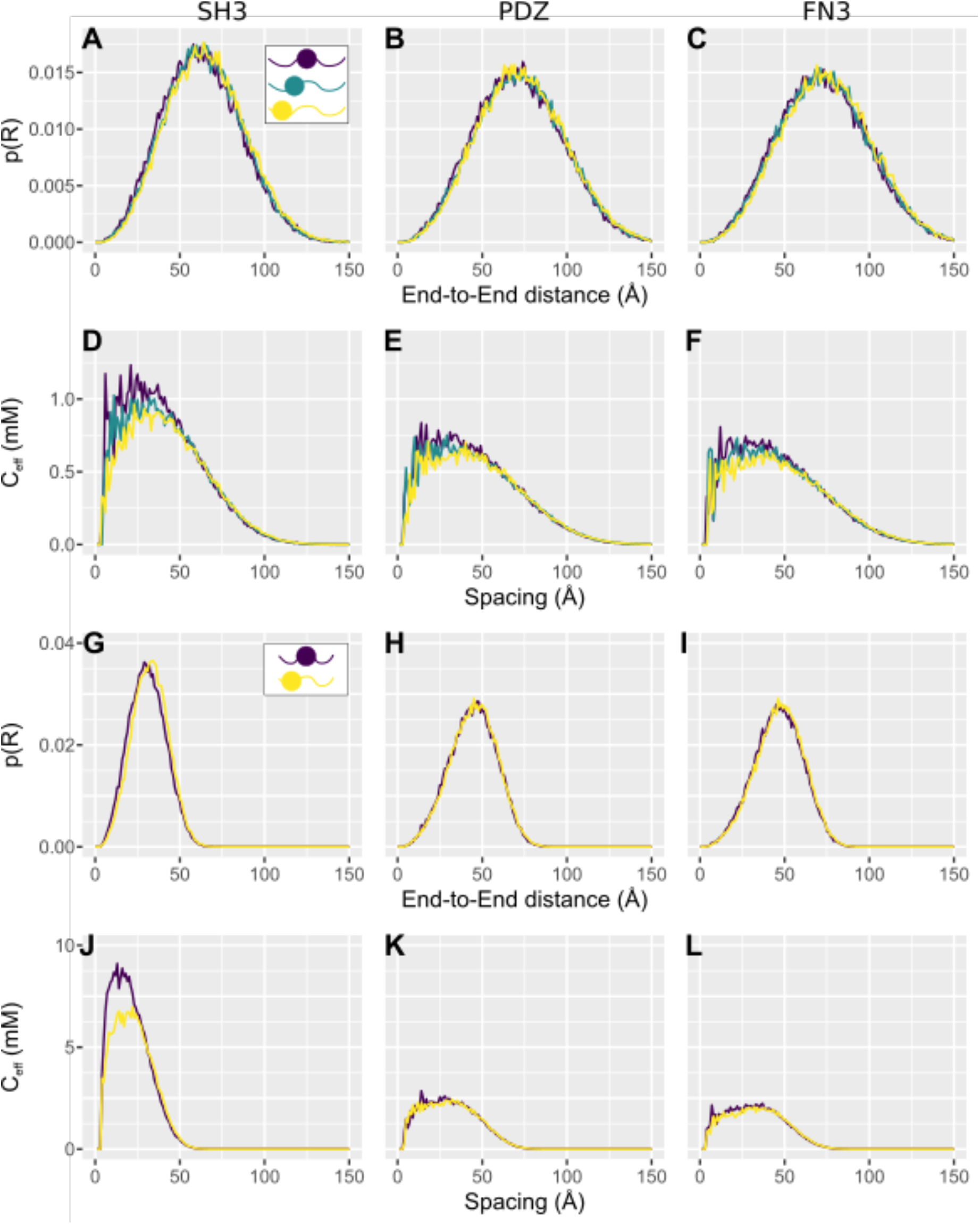
Multiple linkers can mostly be combined into a single longer linker. Predicted end-to-end distance distributions and C_eff_ from 50.000 member ensembles for linkers composed of a folded domain flanked by two disordered segments. (A-F) A total of 100 disordered residues distributed as 50-50, 25-75 and 5-95. (G-L) A total of 30 disordered residues 15-15 and 5-25. Structures used: (A, D, G, J) SH3 domain from (PDB: 1w6x). (B, E, H, K) PDZ3 domain from PSD-95 (PDB: 5d13). (C, F, G, H) 11^th^ FN3 domain from fibronectin (PDB: 5dft).

The ensembles suggest that different combinations of folded and disordered segments have similar *C_eff_* profiles if the total length of the disordered segments are identical. Fig. 4 compares end-to-end distance distributions for a folded domain surrounded by flexible linkers with similar length but distributed differently. Generally, the predicted *C_eff_* was slightly higher when the linker was distributed equally around the folded domain corresponding to a higher sampling of close contacts. The likely cause of this deviation is the steric effect of the folded domain, which affects the proximal region of the linkers most strongly. The deviation in the *C_eff_* was largest for the linker architecture with short flexible segments surrounding the SH3 domain. The SH3 has a short distance between the N- and C-termini (7.1 Å), which likely enhanced the steric effects of the linkers. However, these deviations are minor, and for most practical purposes the end-to-end distance distribution and *C_eff_* profile depend mostly on the total length of flexible segments, and much less on how they distribute around the folded domain.

#### Is it valid to approximate folded domains with a stiff rod?

Next, we sought to test how a folded domain inserted into a long linker affects *C_eff_*. Geometric considerations suggest that a folded domain has two effects: It act as a rigid spacer within the linker, and it adds an excluded volume the flexible segments cannot enter. Previous work suggests that the rigid spacer effect is the dominant effect, and that the domain can often be approximated by a stiff rod with a length corresponding to the distance between the N- and C-termini of the domain (r_NC_).^25^ This simplification ignores other effects of the shape and size of the folded domain. To test the rod approximation, we inserted 20 folded domains (Fig. 5A) that are roughly globular and span a range of sizes and r_NC_ values in the middle of a 100-residue disordered linker (Fig. 5B). To approximate a rod, we built ideal poly-alanine α-helices with r_NC_ in the range from 10-100 Å, and generated ensembles for each rod in the middle of a 100-residue GS-linker. The mean end-to-end distances from the globular domains largely follow the same curve as the helical rods (Fig. 5C) although some of the large domains occupy a slightly more expanded ensemble likely due to the excluded volume of the domain. The effective concentration was calculated at spacings just below (50 Å) and above (100 Å) the mean end-to-end distance in the fully disordered linker (Fig. 5DE). *C_eff_* followed the same trends for the folded domains as the helical rods with a monotonous decrease with increasing r_NC_ at a spacing of 50 Å (Fig. 5D), a maximum around r_NC_ of 60 Å for a spacing of 100 Å (Fig. 5E). This suggested that the effects of a folded domain in a linker can be approximated by a rigid rod and that the bulk of the domain plays a subsidiary role. Furthermore, it also showed that a domain of a fixed size will have a different impact, depending on where the flexible linkers are attached: If the termini are on the same side of the domain the effect will be much smaller than if they are located on the opposite side.

**Figure 5:**
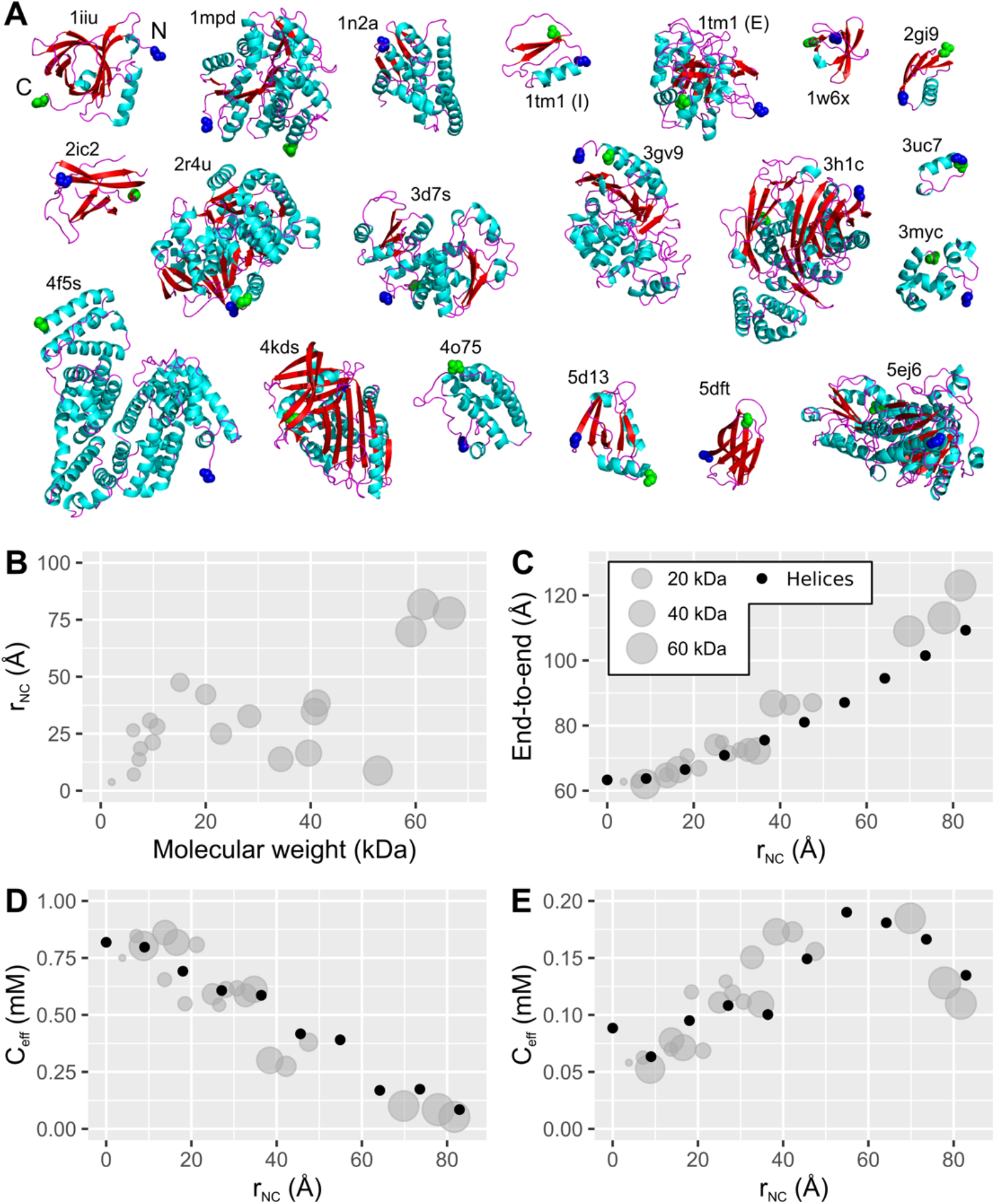
Globular domains in linkers can be approximated by a rigid rod. (A) Structures of the twenty globular domains that were inserted in the middle of a 100-residues GS linker and modelled using EOM. The N-termini (blue) and C-termini (green) are highlighted. (B) Distribution of N-to-C distances of the globular domains (r_NC_) and molecular weights in the folded domains used. (C) The mean end-to-end distance of the linker containing each of the folded domains compared to the rigid α-helices of variable length. The circle area is proportional to the M_W_ of the globular domain. (D and E) *C_eff_* was calculated at spacings of (D) 50 Å and (E) 100 Å. r_NC_ was measured between C_α_ of the first and last residue. The ensemble size was 10.000 for all linkers except those used in Fig. 4 (PDB: 1W6X, 5D13, 5DFT), where the ensemble size was 50.000. The data point plotted at r_NC_ = 0 represents the linker without an inserted domain.

#### The effect of r_NC_

Having validated the rod assumption, we further examined the end-to-end distance distribution and *C_eff_* profile of linker architectures with rods of increasing r_NC_ (Fig. 6A). When r_NC_ is small, the ensemble dimensions are dominated by the disordered chains, but with increasing r_NC_ the dimensions of the rod start to dominate. The increase in end-to-end distance thus accelerates as the length of the rod increases in agreement with (Fig. 5C). These changes mirror a gradual change in the shape of the *C_eff_* scaling profile with insertion of rods of increasing r_NC_ (Fig. 6B). For short rods, *C_eff_* is still highest at small spacings, whereas the profile gradually changes to have a non-zero maximum for longer rods. The distance distribution translates into different dependence of *C_eff_* on the rod length depending on the spacing: At spacings below the average end-to-end distance of the linker without rods, *C_eff_* decreased monotonously with rod length (Fig. 5B). Whereas at long spacings (> ~100 Å), *C_eff_* increased with rod length for the range tested here, although this trend will reverse when the rod is much longer than the spacing.

**Figure 6:**
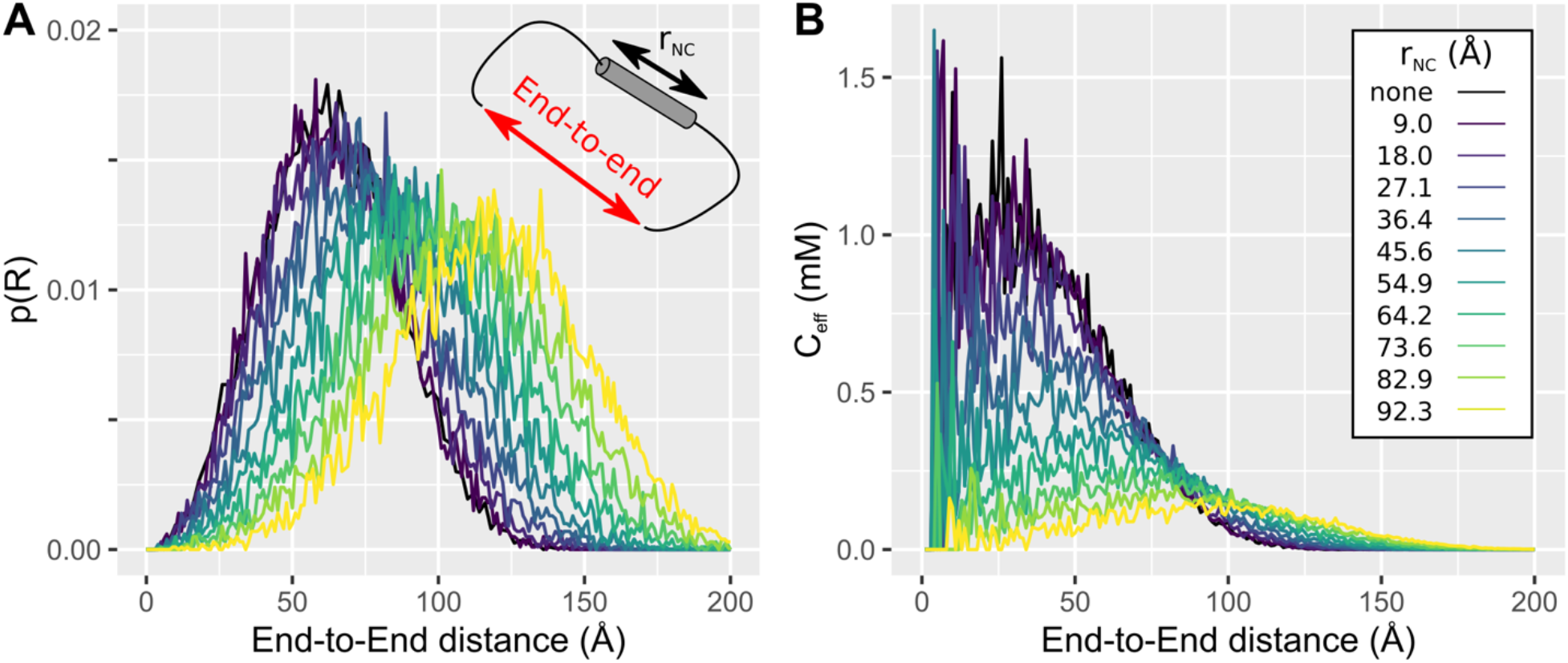
Modelling the effect of folded domain as a stiff α-helix. Rigid poly-alanine α-helices of variable length was inserted in the middle of a 100-residue GS-linker and an ensemble of 10.000 structures was modelled use EOM. For each linker is shown: (A) End-to-end distance distributions, (B) *C_eff_* scaling. The length of the α-helix was increased in steps of 7 residues corresponding roughly to 9 Å steps. The r_NC_ of the rigid rod was measured between the C_α_ of the first and last residues of the helix.

#### Optimal linker architectures for large spacings

Linkers are usually associated with IDRs, but fully disordered linkers span large distances inefficiently. For flexible linkers, the maximum achievable *C_eff_* rapidly drops off with increasing spacing (Fig. 3). The highest achievable *C_eff_* by a disordered linker of optimal length can be calculated using the WLC implemented in “*C_eff_* calculator”.^30^ Using a L_p_ of 6.4 Å to match the “native chain” output of EOM (Fig. 2), the highest achievable *C_eff_* at spacing of 25 Å is 9.6 mM for a 10-residue linker, and 123 μM at 100 Å for a 200-residue linker. To explore whether insertion of a folded domain increases the maximal *C_eff_*, we focused on a binding site separated by a spacing of 100 Å. Linker ensembles were generated with flexible segments flanking α-helical rod corresponding to r_NC_ values of a small (Fig. 7A, 21 residues, r_NC_ = 27.1Å), a medium-sized (Fig. 7B, 42 residues, r_NC_ = 54.9 Å) and a large domain, where the latter has a r_NC_ that approximates the spacing (Fig. 7C, 70 residues, r_NC_ = 92.3 Å). The longest flexible segments considered (200 residues) match the optimal flexible linker and was gradually shortened to reveal the dependence on the length of the flexible segments for each rod (Fig. 7D). We benchmark these values against a fully disordered linker of optimal length (200 residues), which was predicted to have a *C_eff_* of 147 μM from the ensemble at this spacing in reasonable agreement with the WLC (123 μM).

**Figure 7:**
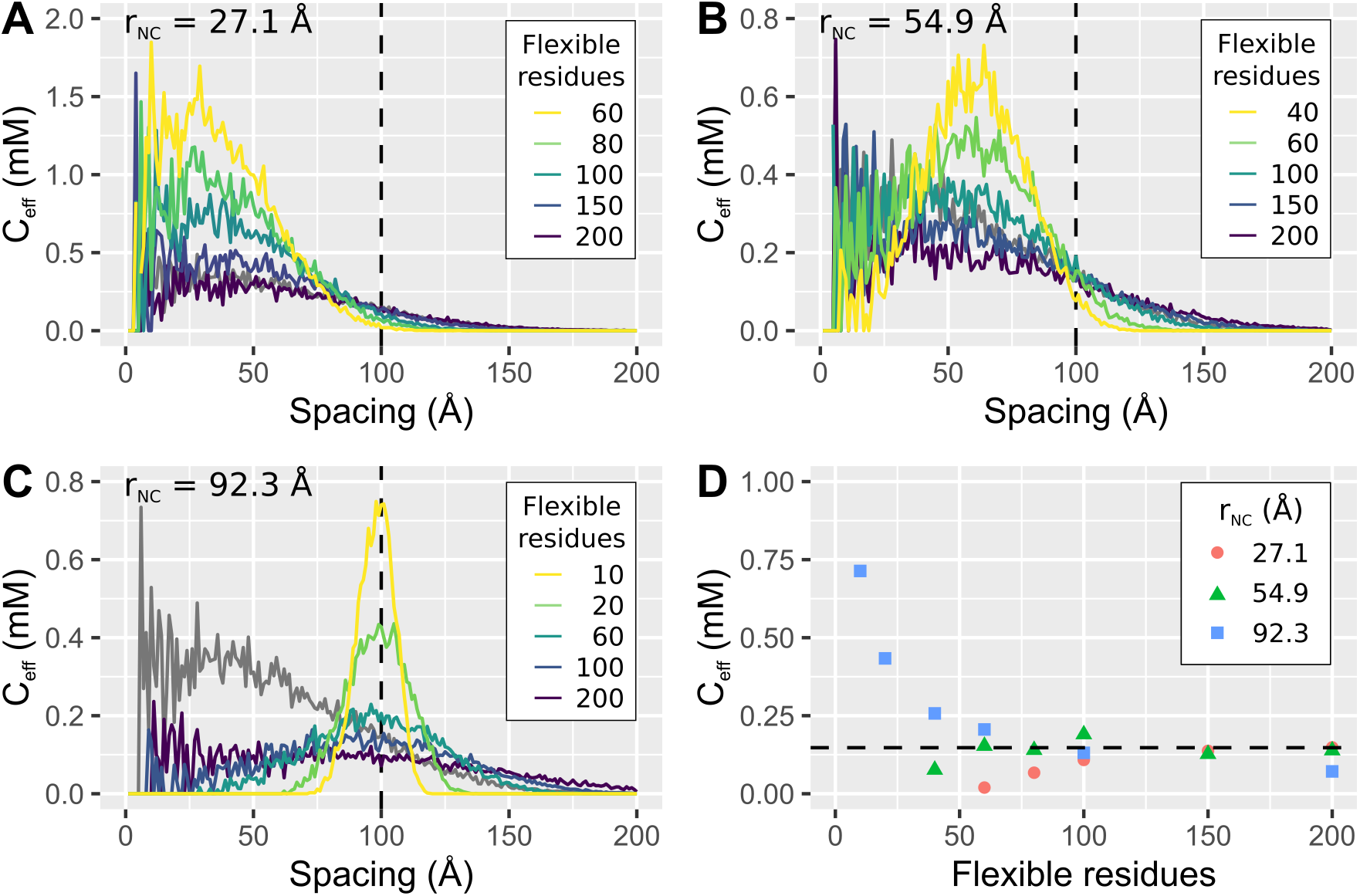
Effective concentrations enforced by linkers composed of a rigid rod flanked by flexible segments of variable length. Ensembles were generated for linkers consisting of flexible segments of variable length with a central α-helix containing (A) 21, (B) 42 and (C) 70 residues corresponding to r_NC_ lengths of 27.1, 54.9 and 92.3 Å. The effective concentrations are compared to the flexible linker that results in the highest C_eff_ at a spacing of 100 Å (200 residues). (D) C_eff_ at a spacing of 100 Å for different linker architectures show that a linker containing a folded domain can exceed the maximal value attainable with a fully flexible linker (147 μM, dashed line), when the r_NC_ of the folded domain matches the spacing between binding sites.

For the short rod, the *C_eff_* profile with 200 flexible residues was similar to the disordered linker and shortening of the flexible segments monotonously decreased *C_eff_* without exceeding the benchmark (Fig. 7D). For the medium length rod, the benchmark *C_eff_* was exceeded slightly when the rod was coupled to 100 flexible residues. However, *C_eff_* was similar to the benchmark for range of flexible segments and did not drop below the benchmark until the linker was shortened to 40 residues (Fig 7D).

The linker architecture containing the longest rod started at low *C_eff_* as it was too extended for the spacing. Shortening of the flexible segments increased *C_eff_* such that it exceeded the benchmark for fewer than ~100 flexible residues and increased noticeably beyond the benchmark for short flexible segments. The origin of this increase can be visualized in the *C_eff_* profile (Fig. 7C). The fully disordered linker spans the large spacing inefficiently because it samples the entire volume between the origin and the binding site. The rigid rod in the linker reduces the sampling of short spacings, and when r_NC_ is matched to the spacing it can increase the highest attainable *C_eff_*. The inclusion of a folded domain into the linker architecture decouples the spacing that needs to be spanned from the volume that is sampled. When r_NC_ is matched to the spacing, shortening of the flanking linkers focus the ligand on the binding site.

For globular proteins, the combinations of a long r_NC_ and short linkers will result in steric clashes not found in the helical model. This suggests that the optimal linker architecture to span large distances involves elongated, rather than globular, folded domains. To test the putative *C_eff_* enhancement of complex linker architectures in a more realistic case, we studied linker architectures containing the rod domain from the scaffolding protein a-actinin. The rod domain is ~250 A long with an anti-parallel heterodimeric coiled-coil architecture (Fig. 8A).^49^ Linker architectures were modelled by inserting disordered segments of variable length at the two C-termini that are separated by 245 Å. Like the α-helical model system, these linker architectures focus the p(r) and *C_eff_* profile at spacings similar to the length of the domain (Fig. 8BC). For comparison at a spacing of 250 Å, the ideal fully disordered linker of 1283 residues achieves a *C_eff_* of ~7 μM. This *C_eff_* is exceeded by a factor of two by a-actinin linkers with a total of 200 disordered residues (Fig. 8C) and rises further as the linkers is shortened. The model in Fig. 8A suggests that this linker architecture would be sterically able to bridge binding sites in most other proteins or assemblies with this spacing. This suggests that the *C_eff_* enhancement from inserting folded domains into disordered linkers will be relevant in many biochemical contexts.

**Fig. 8:**
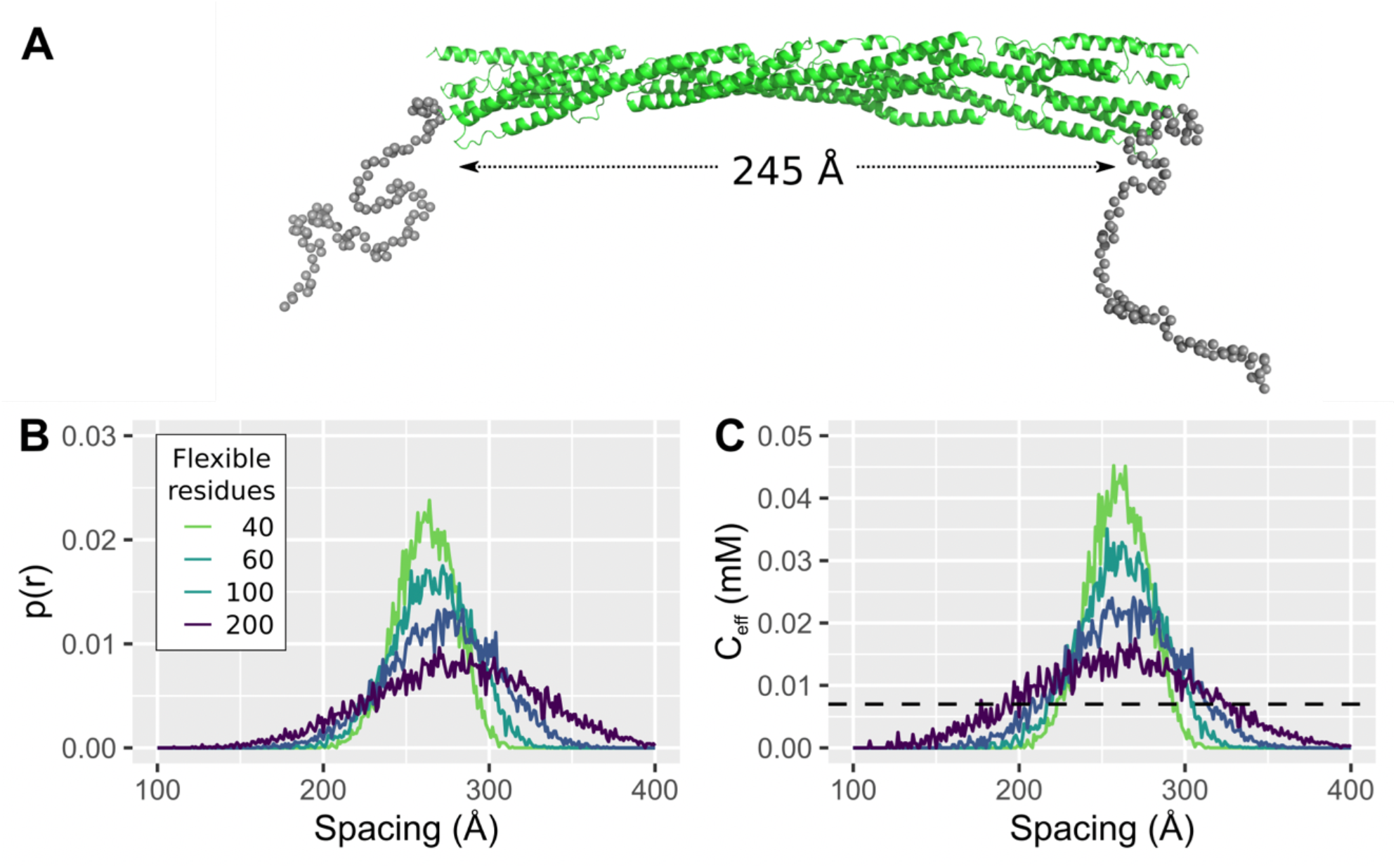
Modelling of linker ensemble of a disordered linkers containing the rod domain from a-actinin. Representative conformation of a linker architecture containing the rod domain of a-actinin (PDB:1HCI)^49^ flanked by 100 flexible residues on either side. The rod domain is an anti-parallel homodimer, so the model was made by appending 100 flexible residues to the C-terminus. The C a-atoms of the C-terminal residues of the two monomers are separated by 245 Å. (B) End-to-end probability distributions (B) and *Cf* profiles (C) of linker architectures containing a-actinin flanked disordered segments of variable length. The dashed line in (C) corresponds to the highest achievable *C_eff_* (7 μM) by a fully disordered linker (1283 residues) at a spacing of 250 Å modelled by WLC with an L_p_ of 6.4 Å.

## Discussion

Here we have described a general method for predicting *C_eff_* enforced by linker architectures from conformational ensembles. Alternatively, such effective concentrations could be calculated from polymer models like the WLC or a measured directly through competition experiments. The method presented here is more cumbersome than the purely analytical models, but is much faster than competition experiments. Where the WLC only describes fully disordered linkers, the ensemble method can determine *C_eff_* from complex linker architectures composed of any combination of flexible and folded segments, and thus expand the scope of linker architectures that can be studied.

*C_eff_* values from ensembles generated with EOM largely match those predicted by the WLC, suggesting that these methods are comparable for fully disordered linkers that can be described by a homopolymer model. An exception occurs at small spacings (Fig. 2D-F), where *C_eff_* values from ensemble simulations are systematically lower, and the *C_eff_* profile has a different shape with a pronounced plateauing. This effect is probably due to the excluded volume, which mainly affects the subset of the ensemble where the ends are close. Since the WLC does not account for excluded volume, the values from the ensemble are likely more accurate.

Predictions can be compared to experimental values from protein biosensors containing interactors joined by a disordered linker. We compare to two biosensors: One that directly measures *C_eff_* through displacement of a tethered peptide, and a Zn(II)-sensor that compares the affinity of a tethered and untethered Zn(II)-dependent interaction.^53^ Both the ensemble and WLC predict *C_eff_* values that are consistently ~3-fold higher than those measured by the *C_eff_* biosensor,^21^ but not the Zn(II)-sensor (Fig. 3).^53^ It is not clear, whether this reflects a real difference between the experiments and the simulations or a systematic error. The experimental *C_eff_* values are derived from a very different approach, and it would be surprising if they were fully comparable to theoretical estimations. For example, the *C_eff_* biosensor contains both the interaction domains and a pair of fluorescent proteins tethered at the end of the linkers. Tethered folded domains increase the excluded volume further^55^ or may interact transiently with the IDR,^56^ which might lead to systematically lower *C_eff_* values compared to the simulations. Furthermore, any structural changes in the binding partners upon binding may change the effective length of the flexible segments and r_NC_ as seen for a two-component biosensor using a different interaction pair, where a different scaling exponent was found for the same linker composition.^13^ Finally, the theoretical estimations assume precisely defined beginnings of the flexible linkers and distances to the binding site. In practice, such estimations will necessarily be approximations as there is a gradual transition from rigid to flexible, and proteins change structure upon binding. Therefore, systematic deviations between experiment and the theoretically predicted values are not surprising and the experiments should thus not necessarily be regarded as a gold standard.

The ensemble used to calculate *C_eff_* could be generated in many ways. The current implementation using EOM has the advantage of simplicity and only relies on open tools. The limitation of this method is that it does not account for sequence-specific interactions, sequence-specific sampling of backbone dihedral angles, or interactions between flexible segments and the folded domains. This is acceptable for a study of the general interplay between rigid and flexible segments of linker architectures but suggests potential future improvements. At the cost of increased computational complexity, sequence-specific behavior could be improved by sampling backbone dihedral angles based on values obtained for e.g. tripeptides.^57^

An alternative strategy would be to determine *C_eff_* from molecular dynamics simulation of the linker architecture. Simulations of intrinsically disordered regions can capture many sequence specific interactions in the IDRs^58^ and between IDRs and folded domains,^55^ however at significantly increased complexity compared to the approach presented here. Recently, *C_eff_* values enforced by a family of disordered linkers were predicted based on simulations.^16^ This study used the average end-to-end distance in the simulation to derive a sequence-specific persistence length, that could then be used to calculate *C_eff_* via the WLC.^59^ This accounts for sequence-specific interactions in the flexible linkers and was critical for developing the idea of “conformational buffering” in linker sequences. Based on the results obtained here, however, it is likely that *C_eff_* can be calculated in a model-free way from the end-to-end distance distribution of the trajectory rather than going via the WLC.

The accuracy of the *C_eff_* determination would be improved if the ensemble was selected or validated using experimental data. The most informative types of experiments include NMR,^60,61^ small angle X-ray scattering,^51^ single-molecule FRET^26^ or any combination thereof.^62,63^ EOM was made for analysis of SAXS data,^31^ and was built on Flexible-Meccano developed for NMR data.^64^ The ensemble generation step can thus directly incorporate refinement of the ensemble using experimental data, but this was not pursued here. Experimental ensembles vary widely in size but are often much smaller than the ~10.000 conformations needed to construct a reliable *C_eff_* profile as seen for the ensembles deposited in the protein ensemble database.^65^ This suggests that for experimental studies aiming at describing tethered reactions a large ensemble should be sought.

Many intra-molecular or intra-complex reactions are tethered by more complex connections than a simple flexible linker. Nevertheless, the term “linker” is often regarded as synonymous with a continuous IDR. We have shown that for the purpose of linking interacting entities, folded domains are easily tolerated in linker architectures, and may even be beneficial compared to a fully disordered linker. This suggests that linker architectures with folded domains embedded could be selected for even in the absence of any other function of the folded domain. This broadens the concept of what can be considered a linker and complicates functional annotations of linkers, which typically presume disorder. The semantic difficulties can be solved by introduction of the concept of linker architecture as a functional designation based on the context in multidomain proteins or complexes rather than an intrinsic property of the sequence.

The rise of multivalent protein drugs such as bivalent antibodies suggests that optimizing linker properties have direct practical and clinical applications. Long disordered linkers sample a large volume, which is disadvantageous when the binding site is placed at a specific spacing. The combination of an elongated folded domain with short flexible linkers thus provides the optimal linker architecture. Similar results were obtained in simulations of a rigid DNA molecule decorated with flexible polyethylene glycol linkers.^24^ Applications that require the proteins to span large distances such as e.g. bivalent binding to antibodies benefit from more rigid linkers such as e.g. a coiled-coil^66^ or (EAAAK)_x_-repeats^67^ that form α-helices.^68^ Alternatively, proline-rich linkers were beneficial in auto-inhibited biosensors as proline increases the rigidity and thus persistence length of the linkers.^69^ Similarly, tandem repeats of small folded domains expand the reach of antibodies more efficiently than flexible linkers.^70^

The *C_eff_* calculated for the linker architectures with a r_NC_ similar to the spacing increases as the linkers shortens, suggesting that the disordered segments are superfluous. This likely demonstrates the limitation of the simplified model used here as the geometry surrounding the binding site is not considered. The IDRs still play a crucial role by providing the conformational flexibility needed to form complexes with minimal physical strain. As illustrated by the case study of the rod domain from a-actinin, coiled-coil domains are an ideal fold to provide a large spacing with a minimum of steric bulk. This may thus explain why long coiled-coil domains are so common in scaffolding proteins.^71^

## Acknowledgements

This work was supported by grants from the Villum Foundation, the Lundbeck Foundation, the Novo Nordisk Foundation and PROMEMO - Center for Proteins in Memory, a Center of Excellence funded by the Danish National Research Foundation (grant number DNRF133). The author thanks Lucia B. Chemes, Nicolas S. González Foutel and Nathalie Wyss for critical feedback on the manuscript.

